# Spiking Recurrent Networks as a Model to Probe Neuronal Timescales Specific to Working Memory

**DOI:** 10.1101/842302

**Authors:** Robert Kim, Terrence J. Sejnowski

## Abstract

Cortical neurons process and integrate information on multiple timescales. In addition, these timescales or temporal receptive fields display functional and hierarchical organization. For instance, areas important for working memory (WM), such as prefrontal cortex, utilize neurons with stable temporal receptive fields and long timescales to support reliable representations of stimuli. Despite of the recent advances in experimental techniques, the underlying mechanisms for the emergence of neuronal timescales long enough to support WM are unclear and challenging to investigate experimentally. Here, we demonstrate that spiking recurrent neural networks (RNNs) designed to perform a WM task reproduce previously observed experimental findings and that these models could be utilized in the future to study how neuronal timescales specific to WM emerge.

## 1 Introduction

Previous studies have shown that higher cortical areas such as prefrontal cortex operate on a long timescale, measured as the spike-count autocorrelation decay constant at rest [1]. These long timescales have been hypothesized to be critical for performing working memory (WM) computations [2, 3], but it is experimentally challenging to probe the underlying circuit mechanisms that lead to stable temporal properties.

Recurrent neural network (RNN) models trained to perform WM tasks could be a useful tool if these models also utilize units with long heterogeneous timescales and capture previous experimental findings. However, such RNN models have not yet been identified. In this study, we construct a spiking RNN model to perform a WM task and compare the emerging timescales with the timescales derived from the prefrontal cortex of rhesus monkeys trained to perform similar WM tasks. We show that both macaque prefrontal cortex and the RNN model utilize units/neurons with long timescales during delay period to sustain stimulus information. In addition, the number of units with long timescales was significantly reduced in the RNN model trained to perform a non-WM task, further supporting the idea that neuronal timescales are task-specific and functionally organized.

## 2 Spiking RNN model

We employed a spiking RNN model based on leaky integrate-and-fire (LIF) units recurrently connected to one another. These units are governed by:

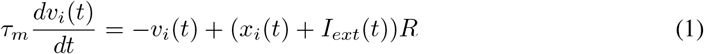

where *τ_m_* is the membrane time constant (10 ms), *v_i_*(*t*) is the membrane voltage of unit *i* at time *t, x_i_* (*t*) is the synaptic input current that unit *i* receives at time *t, I_ext_* is the external input current, and *R* is the leak resistance (set to 1). The synaptic input current (*x*) is modeled using a single-exponential synaptic filter applied to the presynaptic spike trains:

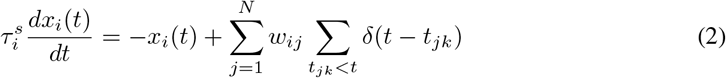

where 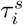 is the synaptic decay time constant of unit *i, w_i,j_* defines the synaptic connectivity strength from unit *j* to unit *i*, and the second summation refers to the spike train produced by unit *j*.

We used the method that we previously developed in [4] to construct LIF RNNs that performed a delayed match-to-sample task (DMS; Figure 1A top). Briefly, we trained several continuous-variable rate RNNs to perform the DMS task using a gradient descent algorithm, and the trained networks were then mapped to LIF networks. In total, we trained 40 RNNs of 200 units (80% excitatory and 20% inhibitory units) to perform the task. The synaptic decay constants (*τ^s^*) were optimized and constrained to vary between 20 ms and 125 ms, but the major findings presented here did not change when the synaptic decay constants were not optimized (i.e. fixed to a constant value; see Section A). All the units from the trained RNNs that satisfied the preprocessing criteria were pooled for the spike-count autocorrelation analysis (see Section 4).

**Figure 1:**
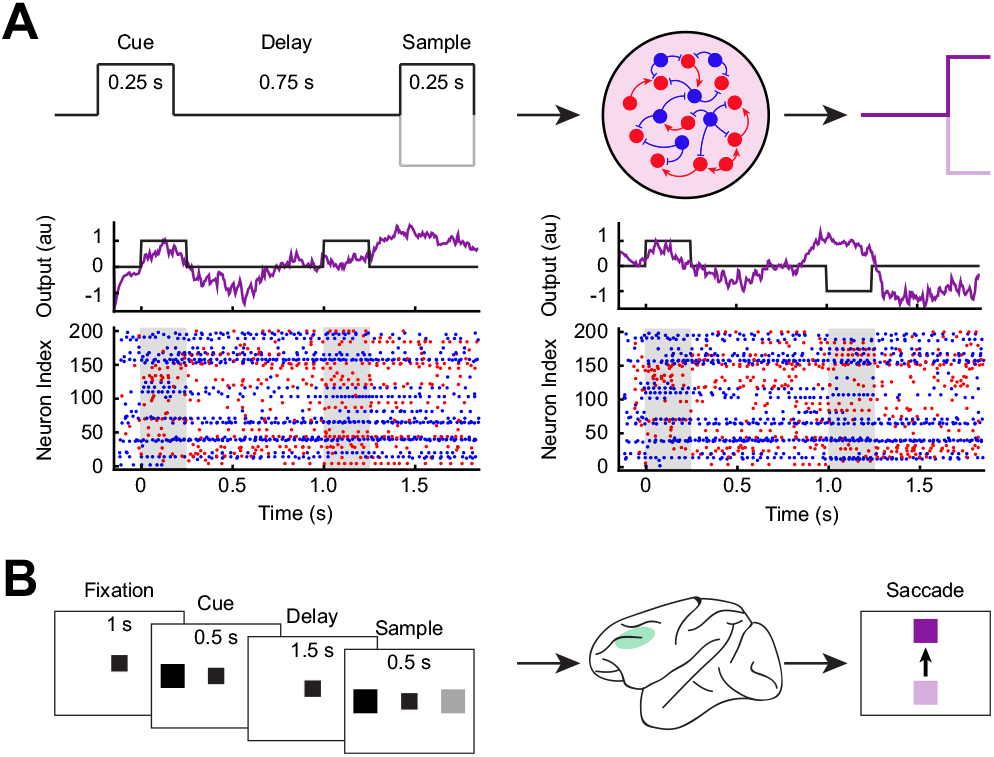
RNN and experimental task paradigms. **A.** DMS task paradigm used to construct the RNN model. Network output and spike rasters from an example RNN model. **B.** DMS task paradigm used by Constantinidis et al. [5] for the experimental data.

The task began with a 1 s of fixation period (i.e. no external input) followed by two sequential input stimuli (each stimulus lasting for 0.25 s) separated by a delay period (0.75 s). The input signal was set to either −1 or +1 during the stimulus window. If the two sequential stimuli had the same sign (−1/-1 or +1/+1), the network was trained to produce an output signal approaching +1 after the offset of the second stimulus (Figure 1A). If the stimuli had opposite signs (−1/+1 or +1/-1), the network produced an output signal approaching −1 (Figure 1A).

## 3 Experimental data

To ensure that the spiking RNN model we employed here is a valid model for investigating neuronal timescales observed in the prefrontal cortex, we compared the findings from our model to the findings obtained from a publicly available dataset [6, 7, 5]. The dataset contains single-neuron spike train recordings from ventral and dorsal prefrontal cortex of four rhesus monkeys performing DMS tasks (Figure 1B). The spike trains of 3257 neurons recorded in the dorsolateral prefrontal cortex (dlPFC) were analyzed for the spike-count autocorrelation analysis. More details regarding the dataset and the tasks can be found in [6, 7].

## 4 Neuronal timescales specific to working memory

### Spike-count autocorrelation decay time constants

To characterize the temporal receptive field, we computed the decay time constant of the spike-count autocorrelation for each unit/neuron during the fixation period [1]. For each unit/neuron, we first binned the spike trains (during the fixation period) over multiple trials using a non-overlapping 50-ms moving window. Since the fixation period duration was 1 s for both RNN and experimental models, this resulted in a [Number of Trials × 20] spike-count matrix for each unit/neuron. For the experimental data, the minimum number of trials required for a neuron to be considered for analysis was 11 trials. The average number of trials from all the neurons included in the analysis was 84.8 ± 34.5 trials. For the RNN model, we generated 50 trials for each unit.

Next, a Pearson’s correlation coefficient (*ρ*) was computed between two time bins (i.e. two columns in the spike-count matrix) separated by a lag (Δ). The coefficient was calculated for all possible pairs with the maximum lag of 650 ms. The coefficients were averaged for each lag value, and an exponential decay function was fitted across the average coefficient values 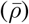 using the Levenberg-Marquardt nonlinear least-squares method:

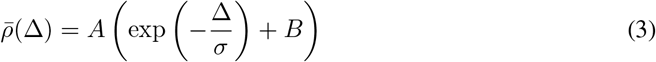

where *A* and *B* are the amplitude and the offset of the fit, respectively. The timescale (*σ*) defines how fast the autocorrelation decays.

The following three inclusion criteria (commonly used in previous experimental studies) were applied to the RNN model and the experimental data: (1) 0 < *σ* ≤ 500 ms, (2) *A* > 0, and (3) a first decrease in *ρ* earlier than Δ = 150 ms. In addition, the fitting was started after a first decrease in autocorrelation.

As shown in Figure 2 (left), the timescales extracted from the untrained RNNs (sparse, random Gaussian connectivity weights; 2769 units from 40 RNNs satisfied the inclusion criteria) were right-skewed. On the other hand, the trained RNNs (841 units from 40 RNNs satisfied the inclusion criteria) and the experimental data (959 units from 4 monkeys satisfied the inclusion criteria) were heavily left-skewed, suggesting that both trained model and data contained predominantly units with long timescales (Figure 2). The distributions and the average autocorrelation values from the RNN model and the experimental data were within those previously reported [1–3].

**Figure 2:**
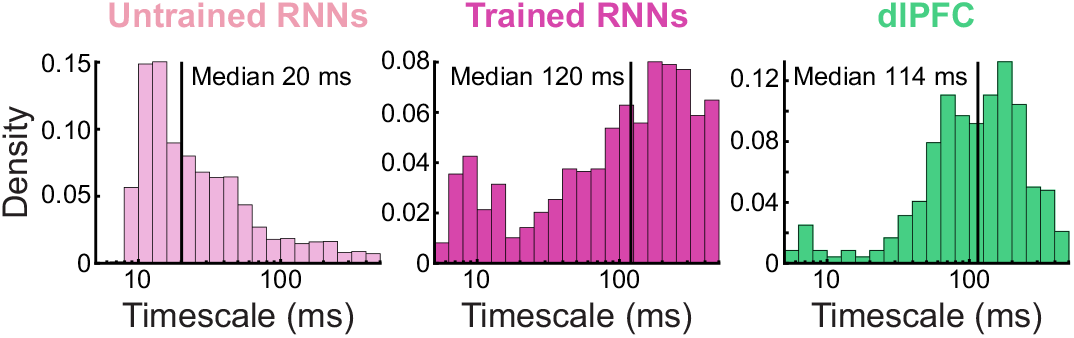
Histograms of the intrinsic timescales extracted from all the units in the RNNs (before and after training) and the experimental data. Solid vertical line, median.

### Units with long neuronal timescales encode information robustly

Next, we investigated to see if the units/neurons with longer timescales were involved with more stable coding using cross-temporal decoding analysis [2]. For each cue stimulus identity, the trials of each neuron were divided into two splits in an interleaved manner (i.e. even vs odd trials). All possible pairwise differences (in instantaneous firing rates) between cue conditions were computed within each split. Finally, a Fisher-transformed Pearson correlation coefficient was computed between the pairwise differences of the first split at time *t*_1_ and the differences of the second split at time *t*_2_. A high Fisher-transformed correlation value (i.e. high discriminability) represents a reliable cue-specific difference present in the network population.

We performed the above analysis on short and long neuronal timescale subgroups from the experimental data and the RNN model. A unit/neuron was assigned to the short *σ* group if its timescale was smaller than the lower quartile value. The upper quartile was used to identify units/neurons with large autocorrelation decay time constants. There were 122 short *σ* and 128 long *σ* neurons for the experimental data. For the RNN model, there were 210 units in each subgroup.

The cross-temporal discriminability matrices (Figure 3) indicate that stronger cue-specific differences across the delay period were present in the long σ subgroup compared to the short σ subgroup for both experimental data and the RNN model. These results are consistent with the previous experimental findings [2, 3], and suggest that longer neuronal timescales correspond to more stable coding.

**Figure 3:**
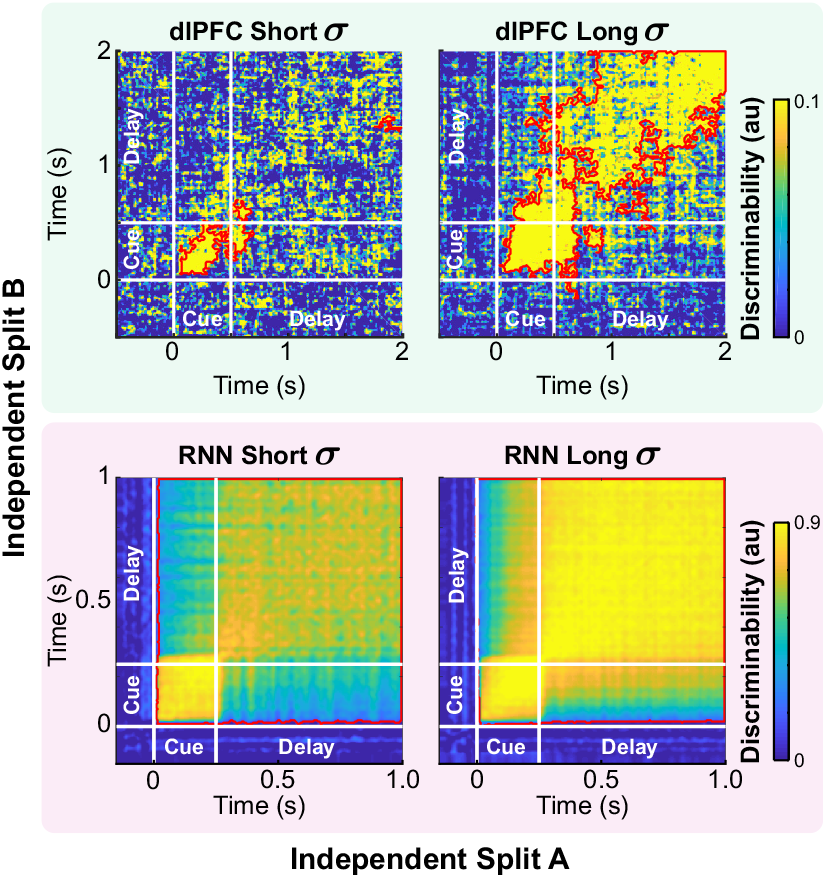
Cross-temporal discriminability scores from the experimental (top) and RNN (bottom) models. Red contours indicate significant discriminability (clusterbased permutation test; *P* < 0:05).

### Task-specific temporal receptive fields

Murray et al. [1] demonstrated that the hierarchical organization of the neuronal timescales from different cortical areas closely tracks the anatomical hierarchical organization. For instance, sensory areas important for detecting incoming stimuli house predominantly neurons with short timescales. On the other hand, higher cortical areas including prefrontal areas may require neurons with stable temporal receptive fields that are capable of encoding and integrating information on a longer timescale.

To investigate if such functional specialization also emerges in our spiking model, we trained another group of spiking RNNs (40 RNNs) on a simpler task that did not require WM. The task paradigm is modeled after a Go-NoGo task, and required the RNNs to respond immediately after the cue stimulus: output approaching −1 for the “−1” cue and +1 for the “+1” cue. Each cue stimulus lasted for 125 ms. This specific task paradigm, which we refer to as Go-NoGo task, was chosen since it does not involve WM, and it has been widely used to study how primary sensory areas process sensory information. Apart from the task paradigm, all the other model parameters were same as the RNNs trained to perform the DMS task.

The autocorrelation decay timescales extracted from the RNNs trained to perform the Go-NoGo task were significantly shorter than the timescales obtained from the working memory RNNs (Figure 4). The RNNs contained fewer units with long timescales compared to the DMS RNNs (Figure 4A), and the timescales averaged by network were also significantly faster than the average timescales from the DMS networks (Figure 4B). In addition, the average autocorrelation timecourse for the Go-NoGo networks decayed faster than the one from the DMS RNNs and resembled the timecourses obtained from the primary somatosensory cortex and the medial-temporal area in the visual cortex (see Figure 1C in [1]). These findings indicate that the neuronal timescales of our RNN models are task-specific and possibly organized in a hierarchical fashion.

**Figure 4:**
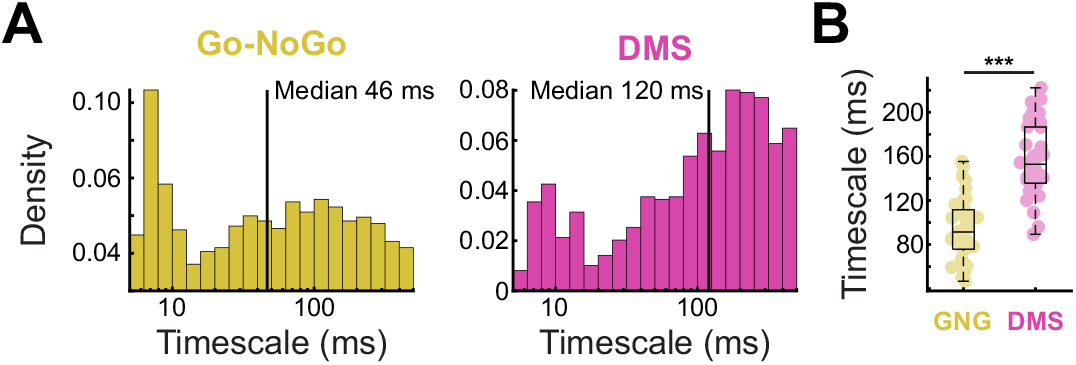
**A.** Neuronal timescale distributions from Go-NoGo and DMS RNN models. 1143 units from the Go-NoGo RNNs; 841 units from the DMS RNNs; Solid vertical line, median. **B.** Average timescales from 40 RNNs for each task model (GNG and DMS). Each circle represents the average value from one RNN. Student’s t-test, \*\*\**P* < 0.0001.

## 5 Conclusions and future work

In this study, we employed a spiking RNN model of WM to investigate if the model exhibits and utilizes heterogeneous timescales for prolonged integration of information. We validated the model using an experimental dataset obtained from rhesus monkeys trained on WM tasks: the model and the primate prefrontal cortex both displayed similar heterogeneous neuronal timescales and incorporated units/neurons with long timescales to maintain stimulus information. The timescales from the RNN model trained on a non-WM task (Go-NoGo task) were markedly shorter, since units with long timescales were not required to support the simple computation. Future works include characterizing the network dynamics and the circuit motifs of the DMS RNN model to elucidate connectivity structures required to give rise to the diverse, stable temporal receptive fields specific to WM.

## Acknowledgments

We are grateful to Ben Tsuda for helpful discussions and feedback. We also thank Jorge Aldana for assistance with computing resources. We also gratefully acknowledge the support of NVIDIA Corporation with the donation of the Quadro P6000 GPU used for this research.

## A Appendix: Effects of synaptic decay constants on neuronal timescales

Here, we demonstrate that the timescales that we obtained from the two RNN models (Go-NoGo and DMS) are not largely driven by the synaptic decay time constants (*τ^s^*) that we optimized to construct the spiking RNNs. For each task model, we trained 40 RNNs with the synaptic decay constants fixed to 125 ms. Even though all the units now had *τ^s^* = 125 ms, the timescale distributions from both models were largely preserved (Figure 1A), and the hierarchy was also maintained (Figure 1B). For the Go-NoGo model, the average timescale values (averaged by network) obtained from the *τ^s^* optimized RNNs were significantly smaller than the timescales computed from the RNNs with *τ^s^* = 125 ms: 93.8 ± 27.6 ms and 109.1 ± 21.0 ms for the *τ^s^* optimized and fixed RNNs, respectively. On the other hand, the average timescales from the *τ^s^* optimized DMS RNNs were significantly larger than the ones extracted from the fixed *τ^s^* networks: 157.0 ± 32.0 ms and 141.1 ± 27.6 ms for the *τ^s^* optimized and fixed RNNs, respectively. Therefore, increasing the synaptic decay time constant for all the units to 125 ms did not necessarily lead to increased neuronal timescales, suggesting that connectivity patterns and structures might play a bigger role in governing task-specific timescales.

**Figure 1:**
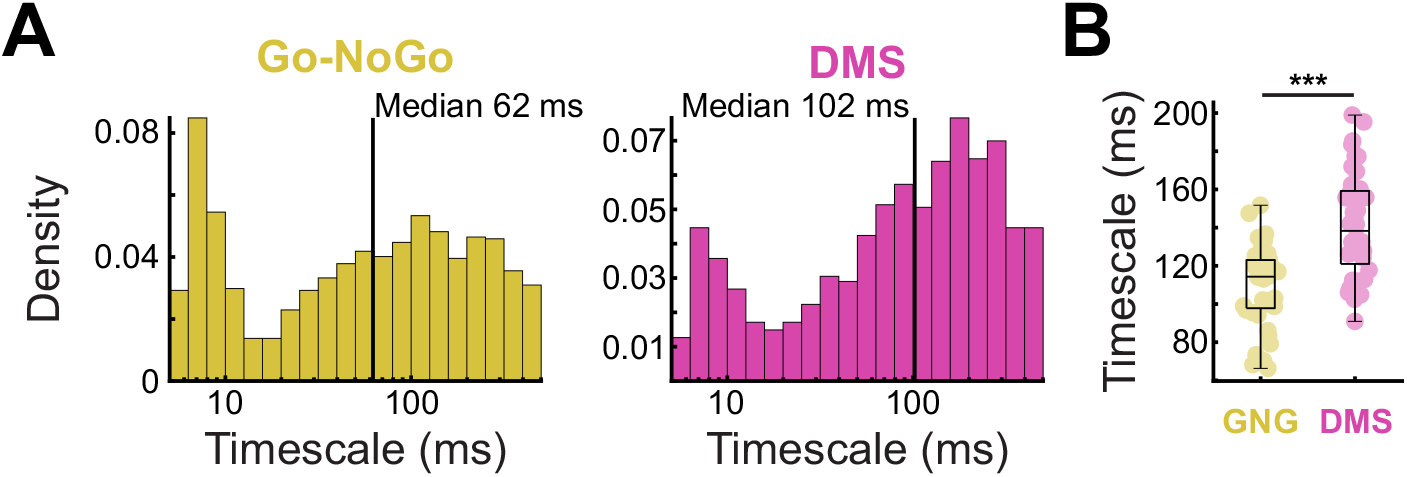
**A.** Neuronal timescale distributions from Go-NoGo and DMS RNN models with *τ^s^* fixed to 125 ms. 1353 units from the Go-NoGo RNNs; 1098 units from the DMS RNNs; Solid vertical line, median. **B.** Average timescales from 40 RNNs for each task model (GNG and DMS). Each circle represents the average value from one RNN. Student’s t-test, \*\*\**P* < 0.0001.

